# Exhaustive identification of genome-wide binding events of transcriptional regulators with ICEBERG

**DOI:** 10.1101/2023.06.29.547050

**Authors:** Anna Nordin, Pierfrancesco Pagella, Gianluca Zambanini, Claudio Cantù

**Affiliations:** Wallenberg Centre for Molecular Medicine, Linköping University, Sweden; Department of Biomedical and Clinical Sciences, Division of Molecular Medicine and Virology; Faculty of Medicine and Health Sciences; Linköping University, Sweden

## Abstract

Genome-wide protein interaction assays aspire to map the complete binding pattern of gene regulators. However, common practice relies on replication and high stringency statistics which favor false negatives over false positives, thereby excluding portions of signal which may represent biologically relevant events. Here, we present ICEBERG (Increased Capture of Enrichment By Exhaustive Replicate aGgregation), an experimental and analytical pipeline that harnesses large numbers of CUT&RUN replicates to discover the full set of binding events and chart the line between false positives and false negatives. We employed ICEBERG to map the full set of H3K4me3-marked regulatory regions and β-catenin targets in human colorectal cancer cells. The ICEBERG datasets allow benchmarking of individual replicates, comparison of the performance of peak calling and replication approaches and expose the arbitrary nature of other strategies to identify reproducible peaks. Instead of a static view of genomic targets, ICEBERG established a spectrum of detection probabilities across the genome for a given factor, underlying the intrinsic dynamicity of its mechanism of action, and permitting to distinguish frequent from rare regulation events. Finally, ICEBERG discovered instances, undetectable with other approaches, that might underlie novel mechanisms of colorectal cancer progression.

## Introduction

Gene regulation is executed by a combinatorial action of DNA-binding proteins which physically associate with the DNA at regulatory sites across the genome to tether chromatin modifying complexes and RNA Polymerase II and start RNA transcription^1,2^. Characterizing the genome-wide binding profile and positioning of all transcriptional regulators is therefore crucial to understand the dynamics of gene expression across development and disease^3^.

Several technologies have been historically employed to map the binding behavior of gene regulators, among which the popular ChIP-seq (chromatin immunoprecipitation followed by sequencing)^4^ and more recently CUT&RUN (Cleavage Under Targets and Release Using Nuclease)^5^. CUT&RUN in particular is emerging as method of choice, for it does not require cross-linking, only necessitates low cell input^6^, can be adapted to target all types of gene regulators^7^, and its variants – such as CUT&Tag – are suitable for single-cell analyses^8^.

These technologies rely on next generation DNA sequencing (NGS) of the genomic sites associated with the targeted protein, and are therefore intrinsically noisy^9^. Peak-calling algorithms have been developed as necessary tools to track real signal over background: among these, commonly used are MACS2^10^ and the recently adapted for the CUT&RUN more sparse reads SEACR^11^. Yet-unsolved genomics complexities combined with technical artifacts, which include excessive read amplification and mapping biases, can produce spurious signal that is computationally removed by subtracting pre-packaged blacklists of problematic regions^12,13^.

Common practice and working standards developed for ChIP-seq demand the use of experimental duplicates to assess biological reproducibility^14^. Yet, widespread recent application of these technologies generated the consensus that more replicates may improve the reliability of peak identification^15^. The focus remains on adopting stringent statistics and peak reproducibility – often employing the majority rule of “trusting” a peak when called in 2 out of 3 experiments^15^ – as rationale for peak calling. Many pipelines exist to attempt to account for replicates during ChIP-seq peak calling, but these still rely on the initial high stringency statistics^16,17^. These practices are meant to exclude noisy background (true negatives) to make sure that no spurious signal is “called” (false positives) often at the cost of excluding real biological events (false negatives)^18^.

As many before us^15,18^, we have noticed that, across our several CUT&RUN experiments, the overlap between replicates was invariably poor. Moreover, almost paradoxically, the attempt of increasing the rate of peak discovery by increasing the number of replicates led to a dramatic drop in size of the overlapping set across replicates, thus forcing us to rely only on a very small number of reproduced binding instances. These, we submit, must be the tip of an iceberg of biologically relevant events of interest, most of which are discarded.

To solve this issue, we developed ICEBERG (Increased Capture of Enrichment By Exhaustive Replicate aGgregation), an experimental and computational procedure that utilizes numerous CUT&RUN replicates to discover the entire set of binding events and consistently exclude false positives. We applied ICEBERG to profile H3K4me3-marked regulatory regions and β-catenin targets in human colorectal cancer (CRC) cells, a context in which Wnt/β-catenin constitutes the main oncogenic driver and is subject of intense investigation in search of druggable mechanisms to treat disease^19–23^. We showed that ICEBERG approaches the discovery of the entire binding profile of H3K4me3 and β-catenin, uncovering previously undetected cancer relevant binding events and revealing that transcriptional modulators, instead of a fixed number of target sites, are associated to a spectrum of detection probabilities across the genome.

## Results

### CUT&RUN assays are characterized by low concordance across replicates

Across the many CUT&RUN experiments we have performed, we have noticed a typically low overlap between replicates that persists regardless of protein target, peak caller choice, or statistical stringency (Figure 1A). For example, of the total peaks called in our datasets targeting the histone modification H3K4me2 in HEK293T, 62.45% of peaks were called in both replicates. When targeting the transcription factor SOX2 in neural stem cells, the overlap was 48.92%, while in CUT&RUN against the co-factor β-catenin the overlap was only 8.85% (Figure 1B, left column). We refer to the overlap as “concordance”. The low concordance leaves the question open whether the remaining 31.55%, 51.08% and 91.15% of called peaks are real peaks or not. We found this to be a persisting problem also when looking across other published datasets for different gene regulators. In 3 examples of histone modifications, H3K4me3, H3K9me3 and H3K27ac, the overlaps were 48.86%, 56.47%, and 32.51%, respectively (Figure 1B, top row). For the transcription factors Ikaros, SALL4, and GATA1 the overlaps were 37.93%, 1.99%, and 42.30% (Figure 1B, middle row). For the co-factors CBP, HDAC1 and HDAC2, 8.40%, 26.24% and 17.03% of peaks were found concordant (Figure 1B, bottom row). These examples were performed independently by different groups, on different models and analyzed with different pipelines, which reinforces that low concordance is a common outcome in the field. The large number of peaks called in the non-overlapping sets are often assessed by a third replicate to recover more peaks: and while using a 2 of 3 approach can increase the number of retrieved peaks, the number of concordant peaks (that is, those called in 3 of 3 datasets) decreases.

**Figure 1.**
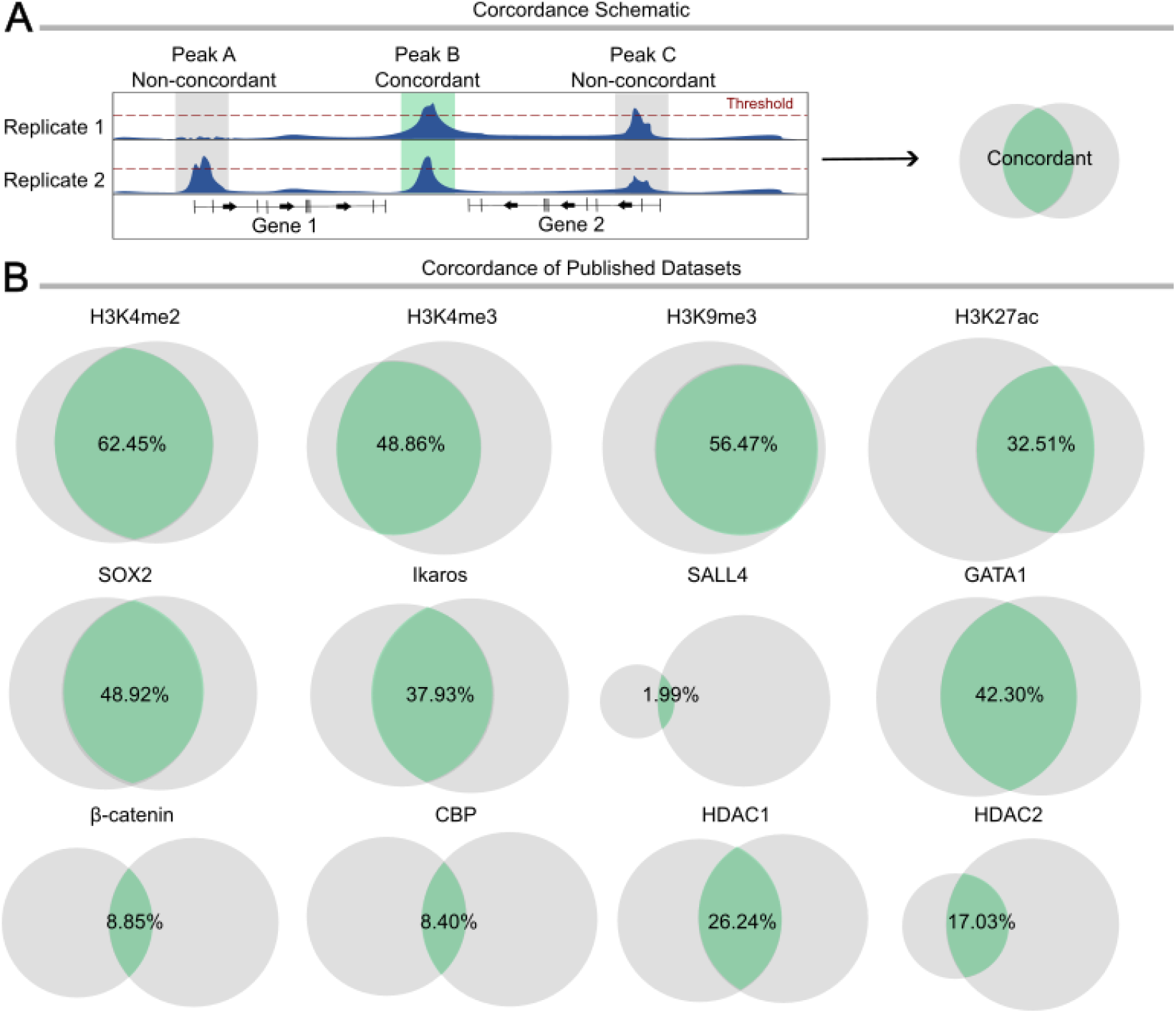
CUT&RUN replicates display low concordance. **A**. Schematic representation of concordance between CUT&RUN replicates, showing peaks present in only one replicate (left), present and called (above red threshold) in both replicates (center) and present in both replicates but only called in one (right). Concordant peaks are represented by the green center of the Venn diagram. **B**. Proportional Venn diagrams showing peak concordance (green sections) across published CUT&RUN datasets for histone modifications (top row), transcription factors (middle row) and other gene regulators (bottom row). Concordant peaks are shown as the percentage of total peaks called.

### Increasing the number of replicates reduces the number of reliable events

To assess how the detection of concordant events changes with increasing replicates, we focused on two targets that, according to analysis above, are characterized by higher and lower concordance. We performed high numbers of replicates of CUT&RUN LoV-U against H3K4me3, β-catenin, and the negative control IgG in human colorectal cancer HCT116 cells over two independent rounds of experiments (Figure 2A). We called peaks in each of the replicates against the IgG negative control with MACS2, using standard threshold settings (q < 0.05), producing individual datasets all of which appear reliable as they display 1) high signal to noise ratio, and 2) enrichment on positive control loci, such as promoters for H3K4me3 and Wnt responsive elements for β-catenin (Figure 2B). Each dataset, when taken alone, would be considered as a reliable genome-wide binding pattern, though together they range in efficiency and number of peaks called. These datasets are not fully concordant, and the low overlap is exacerbated by increasing the number of replicates from 2 to 5 (Figure 2C, Venn diagrams). The strictly concordant peaks (present in all replicates) for both H3K4me3 and β-catenin steadily decrease with increasing replicates. Using the majority approach (2 of 3, 3 of 4, 3 of 5) for H3K4me3 mildly increases the number of considered peaks, while the difference is greater for β-catenin (Figure 2C, bottom graphs).

**Figure 2.**
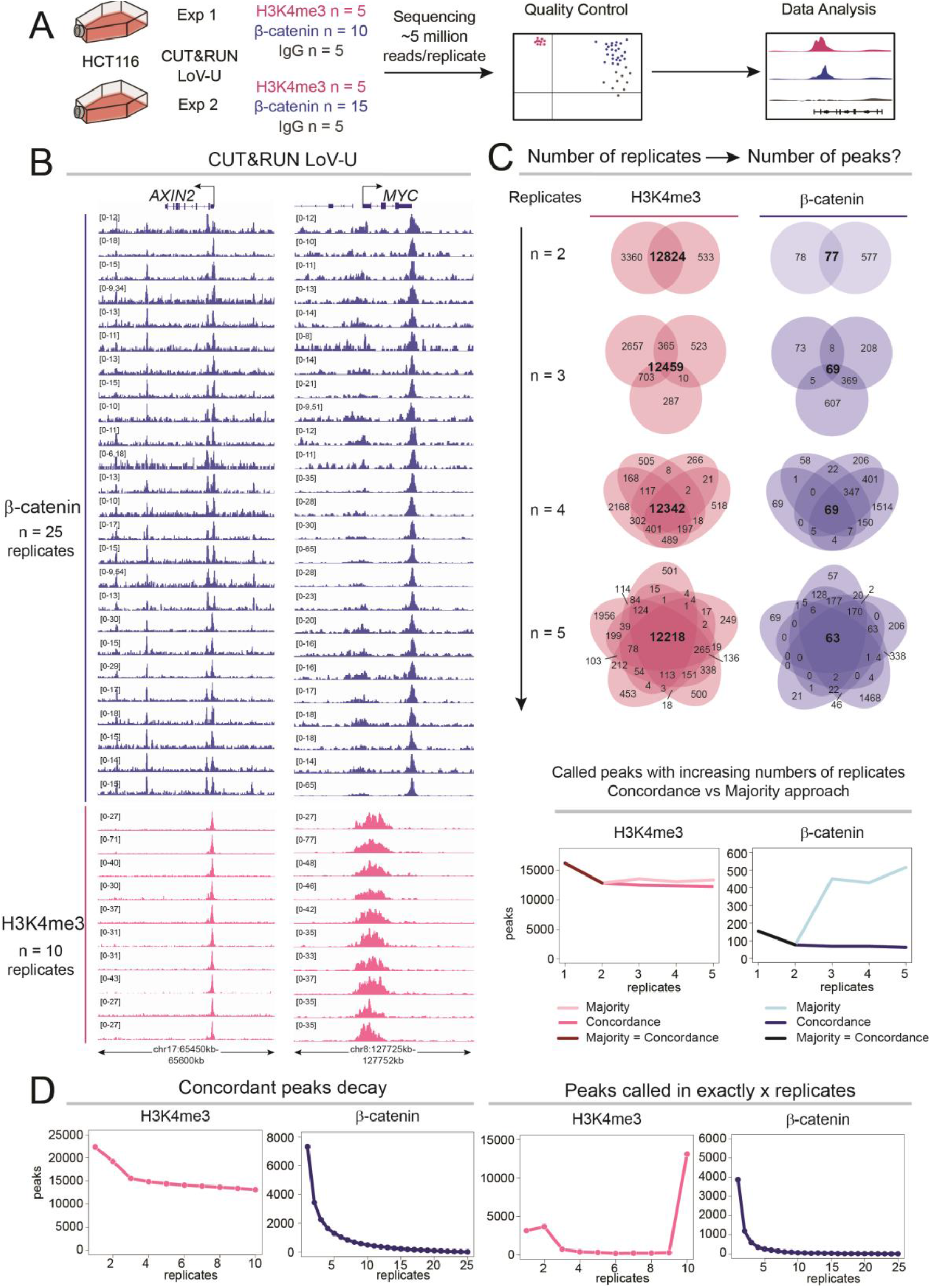
Replication decreases the number of concordant peaks. **A**. Schematic representation of the experimental design. Two rounds of CUT&RUN LoV-U were performed targeting β-catenin (n = 25), H3K4me3 (n = 10) and IgG (n = 10). **B**. Tracks of all performed β-catenin and H3K4me3 replicates at the positive control *AXIN2* and *MYC* loci. All replicates show enrichment and high signal to noise ratio. **C**. Top: Venn diagrams representing concordant peaks (MACS2 q < 0.05). Bottom: Line graphs of the number of concordant and majority approach peaks. **D**. Left: decay curves based on the number of peaks called in at least *x* replicates. The curve of β-catenin displays faster decay than H3K4me3. Right: decay curves based on the number of peaks called in exactly *x* datasets, showing that there are large numbers of peaks only called in subsets of replicates, especially for β-catenin where > 90 % of statistically significant events (q < 0.05) occur in 5 or less replicates.

This analysis raises the question of whether the decreasing concordance would ultimately stop and converge on a small set of the highest confidence binding events. To address this, we leveraged the large number of replicates (25 for β-catenin, 10 for H3K4me3) to build a “decay curve” plotting all peaks called in at least 1, at least 2, at least 3, and so on to finally identify the peaks concordant among all replicates (Figure 2D, left graphs). The H3K4me3 replicates display higher concordance, and this is reflected in a slower decay measured as the lower slope of the curve. On the other hand, the lower concordance of β-catenin replicates is reflected by a rapid initial decay: while 7314 unique peaks are called in at least one replicate, only 13 of these remain concordant in all 25 datasets. This underscores that a relatively large number of events only occur in subsets of replicates and might be therefore rare in their nature. To explore this, we estimated the frequency of their detection by ranking them based on the number of replicates in which they are called, namely plotting those occurring in *only* 1, only 2, only 3 and so on (Figure 2D, right graphs). For H3K4me3, almost all of the peaks were concordant and called in all 10 replicates, and thus likely common events in the cell population, consistent with the known stability of chromatin features^24^. On the other hand, this behavior was different for β-catenin, as > 85% of statistically significant events (q < 0.05) occur in 5 or fewer replicates.

This analysis exposes several analytical problems. For instance, if only 2 or 3 replicates are performed, most binding events of transcriptional regulators risk being missed. Moreover, our analysis imposes the rejection of the majority rule (13 out of 25), which would result in losing more than 90% of detected events for β-catenin. Overall, our data clearly exposes the arbitrary nature of any choice as to what is considered “reproducible”.

### ICEBERG retrieves the complete set of genome-wide binding events

We set out to identify the full set of CUT&RUN peaks for our two targets, H3K4me3 and β-catenin. We constructed an analytical pipeline that would include the information retrieved in all replicates performed: Increased Capture of Enrichment By Exhaustive Replicate aGgregation (ICEBERG). ICEBERG pools equal numbers of random reads from each replicate to build a unique track, where non-specific signals would average into a homogenous background, abrogating false positives, and at the same time allowing the capture of all true signal enrichment, thereby not tolerating false negatives (Figure 3A). ICEBERG entails the repetitive construction of aggregate datasets a total of three times with different read splits from the replicates to minimize the bias due to random read assignment. ICEBERG then employs MACS2 (q < 0.05) to call peaks based on the majority rule (2 out of 3 computational aggregates) against the compilation of the IgG controls (n = 10) and filters the peaks to retain only those present in at least one of the individual experimental replicates at lower stringency (p < 0.01) (Figure 3B).

**Figure 3.**
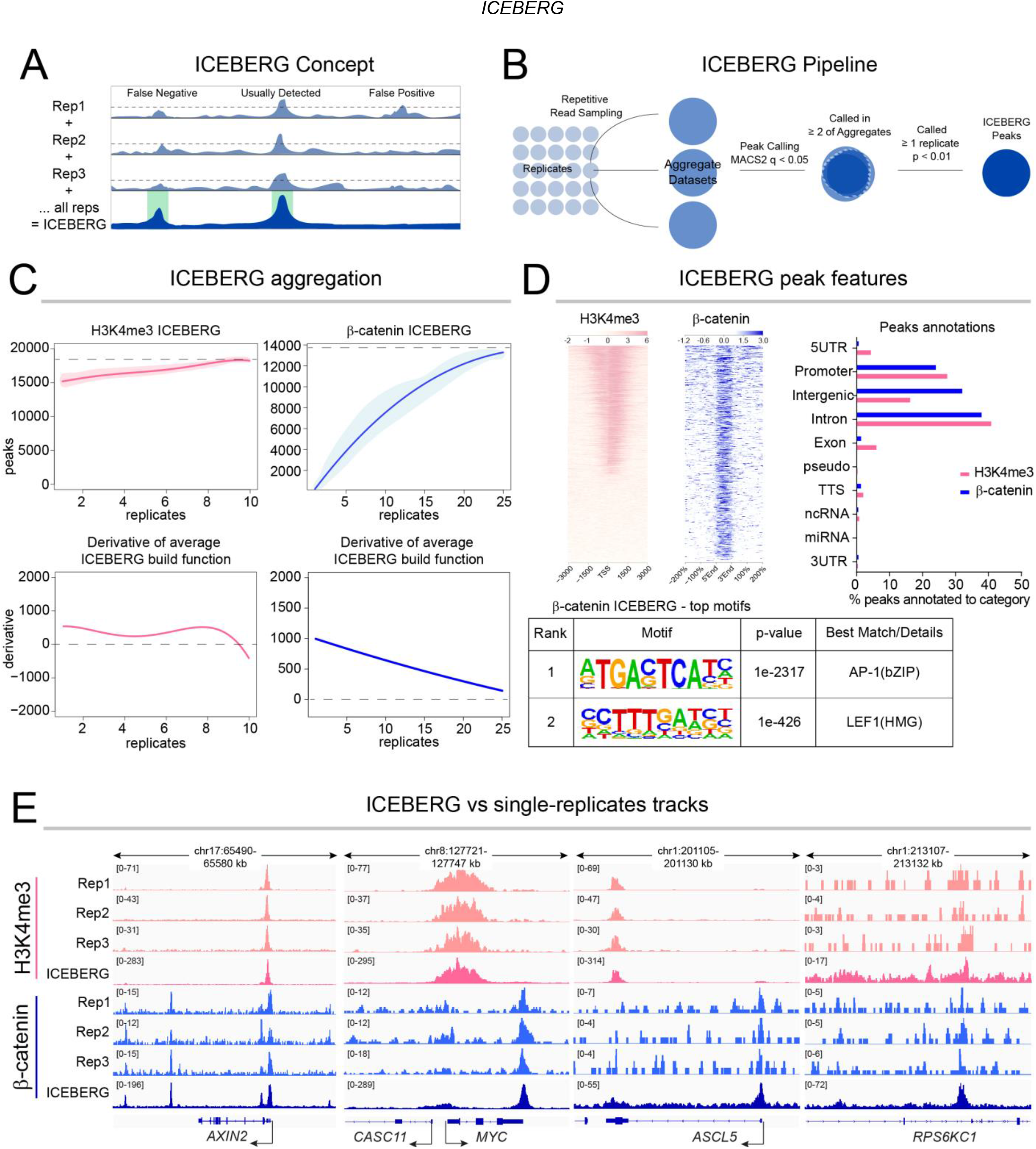
Increased Capture of Enrichment By Exhaustive Replicate aGgregation (ICEBERG). **A**. Schematic of the ICEBERG concept, where the summation of replicates leads to the abrogation of both false negatives and false positives. **B**. Schematic of the ICEBERG pipeline. Reads are sampled randomly three times from all replicates evenly and pooled to form aggregate datasets, which are then peak called and overlapped. Peaks called in at least 2 of 3 aggregates, and also called in at least 1 individual replicate at p < 0.01 are retained. **C**. Top: Regression curve describing the number of called peaks per each added replicate during the ICEBERG aggregate generation. Bottom: Plots of the first derivates of the ICEBERG regression curves. Dashed lines indicate where the derivative equals 0. **D**. Left: Signal intensity plots of the ICEBERG datasets around the transcriptional start sites (TSS) (H3K4me3, left) and peak regions (β-catenin, right), showing high signal to noise ratio. Right: Genomic region annotation of ICEBERG peaks. Bottom: *de novo* motif enrichment in the β-catenin ICEBERG peaks, which displays strong enrichment for the known β-catenin co-regulators AP1 and TCF/LEF. **E**. Example comparisons of the ICEBERG datasets versus individual replicates on the common target loci *AXIN2* and *MYC*, and the previously unknown as targets *ASCL5* and *RPS6KC1*.

In ICEBERG, subsequent aggregation of each individual dataset increases the number of discovered binding events. We fit a polynomial regression function to model the rate of peak discovery with increasing replicates for both H3K4me3 and β-catenin (r^2^ > 0.98) (Figure 3C, top graphs). The shape of the curve is indicative of the rate of discovery. The slope is initially steeper, indicating that each individual replicate adds substantial new information. At higher numbers of replicates, the slope decreases, and the curve begins to plateau, as evidenced by the derivative approaching zero (Figure 3C, bottom graphs). We hypothesize that ICEBERG would allow us to estimate how close we are to the discovery of the full set of binding events of a given factor. For H3K4me3, the regression function reaches a derivative of 0 between 9 and 10 replicates, suggesting ICEBERG may have been performed to exhaustion, identifying a total of 17,641 peaks. For β-catenin, the slope significantly decreases beyond 20 replicates, and the curve is predicted to plateau between 27 and 30 replicates with a total of 13,585 to 13,542 peaks, respectively, which would mean that our 25 replicates with 12,672 peaks have discovered over 93% of the entire β-catenin binding profile.

Several observations support the reliability of the ICEBERG datasets. First, both are characterized by high signal enrichment over the control, which in the case of H3K4me3 conforms to the expected shape of signal around the transcription start sites (Figure 3D, left). Second, genomic annotation of peak positions is comparable with several existing datasets for β-catenin and other Wnt regulators^25,26^ (Figure 3D, right). Third, motif analysis displays known transcriptional coregulators of β-catenin as primary consensus signatures^27^ (Figure 3D, bottom). Signal enrichment can be seen on known targets, as well as on many newly discovered regions (Figure 3E).

### ICEBERG allows comparing of replicates and benchmarking of peak calling strategies

We reasoned that the ICEBERG datasets can be used as a gold-standard against which to benchmark individual CUT&RUN datasets. We decided to focus on two parameters: the percentage of fragments within peaks (FrIP), a typical measurement to determine the efficiency of an individual experiment, and the signal profile around the peak summit, that exemplifies the signal to noise ratio. The ICEBERG datasets allow us to calculate the percentage of fragments of each replicate that fall within the ICEBERG peaks (I-FrIP). The I-FrIP is therefore a better measurement for the efficiency of each replicate. The average I-FrIP for H3K4me3 is 25.06 % (+/-4.34 %) and for β-catenin is 1.50 % (+/-0.46 %) as represented in the violin plots (Figure 4A, left plots for each factor). The difference in I-FrIPs between the two targets is in line with the previously reported varying difficulty in detecting them^7^. Moreover, the violins appear to be inverted: while for H3K4me3 most of the replicates fall within the higher end of the spectrum, indicating higher efficiency is more common, for β-catenin the outlier is instead the most efficient dataset while the majority are less efficient, confirming the intrinsic difficulty of its detection. Similarly, the average signal per million reads varies greatly for both factors (> 2-fold for H3K4me3, > 3-fold for β-catenin), though the profile shape is consistent between replicates (Figure 4A, right plots for each factor). The variability measured by these two metrics is striking as most of the experiments performed could be considered technical replicates and were processed in parallel. This indicates that very small differences between samples can result in drastically different datasets, and that this is likely not reflective of biologically meaningful differences. Therefore, ICEBERG implies that changes in signal intensity from one experimental group to another should be rigorously validated before being considered biologically driven.

**Figure 4.**
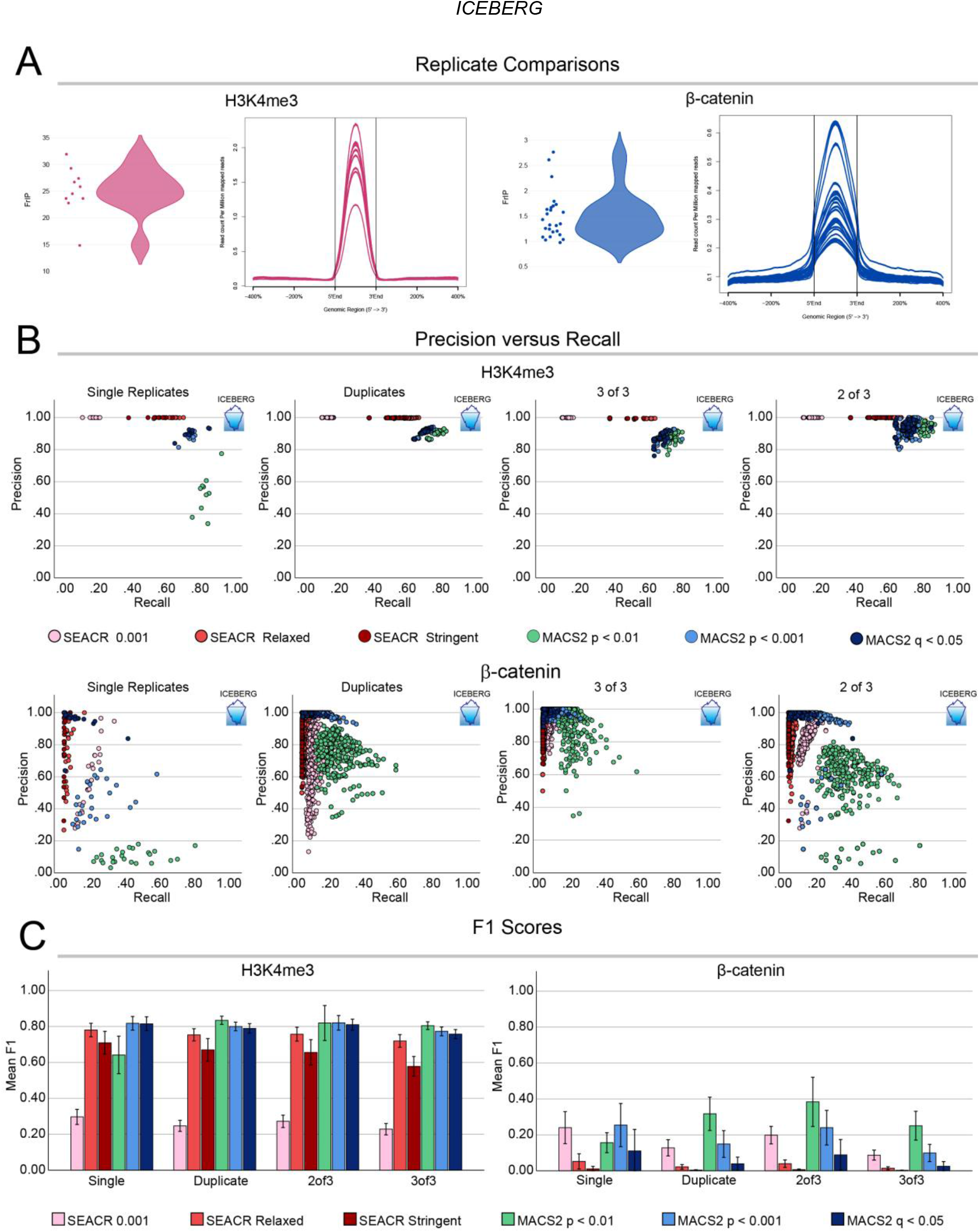
Benchmarking peak calling and replicate strategies using ICEBERG. **A**. Violin plots of I-FrIP scores (fragments within ICEBERG peaks) and signal intensity profiles within ICEBERG peaks of all replicates of H3K4me3 (left) and β-catenin (right). **B**. Graphs showing precision (y-axis, defined as number of peaks called in ICEBERG/total peaks called) and recall (x-axis, defined as number of ICEBERG peaks called/total ICEBERG peaks) of peaks called for each replicate or replicate combination. Different peak calling strategies are shown as different colored dots, and different replicate strategies in different graphs (from left to right: single replicates, duplicates 2 of 2, triplicates 3 of 3, triplicates 2 of 3). H3K4me3 is shown in the top row and β-catenin in the bottom row. In general, H3K4me3 datasets show higher precision and recall, and lower inter-replicate variability than β-catenin datasets, which suffer from lack of recall and have large distributions due to higher variability. **C**. Bar graphs depicting mean F1 scores (harmonic mean of precision and recall) for each replicate and peak calling strategy for H3K4me3 (left) and βcatenin (right). Error bars represent +/-standard deviation.

We leveraged ICEBERG to benchmark the performance of commonly used peak calling strategies done on individual replicates by comparing their precision (fraction of called peaks that fall within the ICEBERG peaks) and recall (fraction of total ICEBERG peaks identified). A high precision indicates a low rate of false positives, whereas a high recall indicates a low rate of false negatives. We called peaks on each replicate using SEACR and MACS2 in various modalities: 1) SEACR on relaxed mode against IgG, 2) SEACR stringent mode against IgG, 3) SEACR 0.001 signal threshold with IgG peaks removed after peak calling, 4) MACS2 against IgG p < 0.01, 5) MACS2 p < 0.001, 6) MACS2 q < 0.05, and plotted their precision and recall. The ICEBERG is by definition situated in the top right corner (precision = 1, recall = 1), whereas the bottom left corner would represent high false positive and high false negative rates, the least desirable outcomes (precision = 0, recall = 0). For single replicates of H3K4me3 the replicates were relatively consistent within each peak calling strategy, all of which except for MACS2 p < 0.01 resulted in a > 0.8 precision. MACS2 p < 0.001 scored the highest recall, closely followed by MACS2 q < 0.05 (Figure 4B, top left). For β-catenin the only strategy with a precision of > 0.8 was MACS2 q < 0.05, though the recall was extremely low at less than 0.2 (Figure 4B, bottom left). Independently of the peak calling strategy used, β-catenin showed high differences between individual replicates, confirming the larger inter-replicate variability when compared with H3K4me3.

### How to approach the ICEBERG with two or three replicates

We have shown that to capture more than 90 % of β-catenin binding events, > 20 datasets are needed. As most CUT&RUN studies are based on a limited number of replicates, we set out to test how well conducting 3 or less than 3 replicates could capture the totality of binding events. We could test this by taking 2 or 3 replicates, overlapping them, and comparing how well in terms of precision and recall they identified the ICEBERG peaks. Due to the variability between replicates (Figure 4A and 4B), the outcome would change based on which combination of samples was chosen. Therefore, we randomly paired our individual datasets in all possible combinations to create duplicates, and in > 50 random combinations to generate triplicates. We overlapped the peaks (2 of 2, 2 of 3, 3 of 3) and plotted the precision and recall scores. Depending on the combination of replicates, the precision and recall varied, but clear distributions could be seen based on the peak calling strategy used. For H3K4me3, precision was high across the board even with the lowest stringency thresholds (Figure 4B, top row). More conservative approaches, like 2 of 2 or 3 of 3, resulted in lower recall than for 2 of 3, where with p < 0.01 it was possible to achieve > 0.9 precision and recall. For β-catenin, higher precision was achieved with replicates versus single datasets, but the recall remained generally low (Figure 4B, bottom row). The least stringent MACS2 p < 0.01 in 2 of 3 could recall up to 60 - 80 % of ICEBERG peaks, but only with up to 0.6 precision. Common practice that favors false negatives over false positives would lead to choosing more stringent approaches at the expense of recall. For an unbiased metric, we calculated F1 scores, which relies on the harmonic mean of precision and recall and provides a balance of false positives and false negatives. According to the highest mean F1 score, the best approach for H3K4me3 would be duplicates with MACS2 p < 0.01, and 2 of 3 MACS2 p < 0.01 for β-catenin (Figure 4C). We suggest that scientists take this information into account when analyzing their data, and choose a strategy based on the biological questions being asked.

### ICEBERG discovers that peaks are distributed in a spectrum of probability of detection

We have shown that ICEBERG approaches the full set of genome-wide binding events, and that most single replicates have an incomplete recall and/or precision. This could be due to two explanations: 1) each replicate identifies a random fraction of the complete set of binding events, and no peaks are preferentially detected across replicates, 2) there could be different types of binding events, each associated with a probability of detection, reflecting their frequency of occurrence in a cell population. To distinguish between these two possibilities, we plotted how many times each ICEBERG peak is identified across replicates (MACS2 q < 0.05). For H3K4me3, each replicate does well in identifying the ICEBERG peaks, as shown by a low slope of the decay curve (Figure 5A, left). The slope of the β-catenin decay curve on the other hand is steep, indicating most ICEBERG peaks are only identified in a few replicates (Figure 5A, right). These patterns can be translated into probabilities of detection by ranking each ICEBERG peak by the exact number of individual replicates in which it is detected.

**Figure 5.**
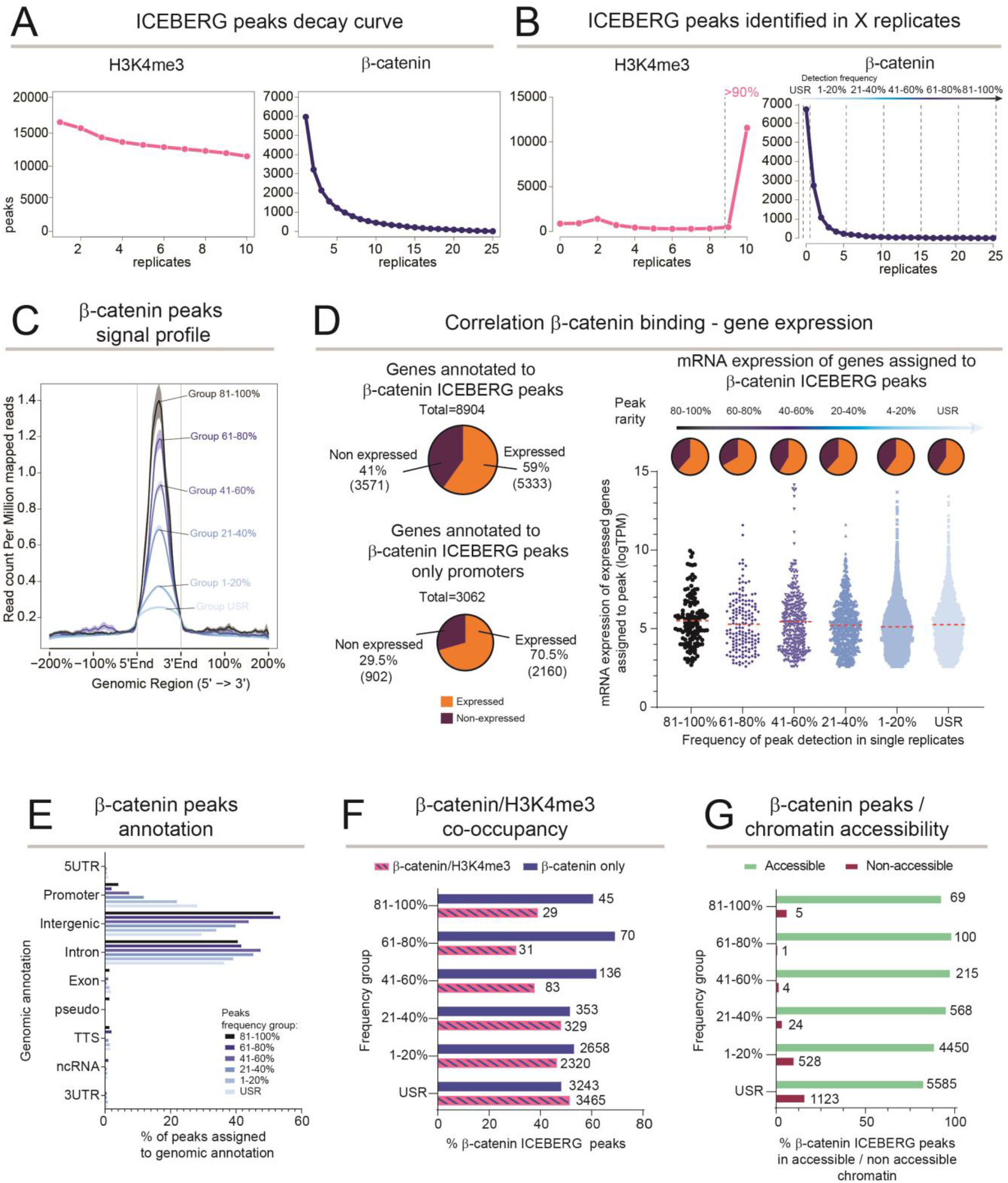
Probability of peak detection is connected to biological function. **A**. Line graphs showing decay of ICEBERG peaks within individual replicates when called with MACS2 q < 0.05. **B**. Line graphs showing ICEBERG peaks called in exactly *x* replicates. The β-catenin ICEBERG peaks are divided into 6 groups based on their probability of detection. **C**. Average signal profiles of different groups of β-catenin ICEBERG peaks. Groups with a higher probability of detection show higher enrichment and a more defined peak summit than less probable peaks. **D**. Left: pie charts showing the number of β-catenin peak-annotated genes (annotated by GREAT on default settings), and promoter associated genes (annotated by HOMER) that are expressed or not expressed according to RNA-seq data (> 5 TPM). Both are higher than expected by chance according to a hypergeometric test. Right: pie charts and graphs showing the number of expressed and non-expressed genes in each group of β-catenin peaks, as well as their log-transformed TPMs. There is no correlation between number of expressed genes or expression levels and increasing β-catenin peak probability. **E**. Bar graph showing genomic region annotation of β-catenin probability groups. With increasing probability of detection, fewer peaks are found on promoters and more within intergenic regions. **F**. Bar graph showing percentage of β-catenin peaks overlapping with H3K4me3 peaks within each group. Similar to the decrease in promoter occupancy, overlap with H3K4me3 decreases slightly with increasing probability. Numbers next to the bar indicate number of events, x-axis represents percentage. **G**. Bar graph of β-catenin peak overlap with ENCODE HCT116 ATAC-seq peaks. The least probable peaks show more overlap with non-accessible chromatin regions versus highly probable peak groups. Numbers next to the bar indicate number of events, x-axis represents percentage.

Over 90% of the H3K4me3 ICEBERG peaks are called in 9 or 10 of the single replicates (Figure 5B, left), further supporting that H3K4me3 is likely easy to detect, homogenous among cells, and a stable feature of the chromatin in HCT116. Almost opposite to this behavior, the majority of β-catenin ICEBERG peaks are rarely detected in single replicates (Figure 5B, right). To explore this, we grouped the β-catenin ICEBERG peaks depending on their probability of detection: undetectable in single replicates at q < 0.05 (USR), called 1 – 5 times (up to 20% probability), called 6 – 10 times (21 – 40%), called 11 – 15 times (41 – 60%), called 16 – 20 times (61 – 80%), and those called 21 – 25 times (81 – 100%). Motif analysis performed individually on each group showed TCF/LEF as a top enriched motif (*de novo* discovery with p < 1e^-10^ for all), confirming that each group, even the ones containing the rarest peaks, consisted of *bona fide* β-catenin binding events. The probability groups were associated with a remarkably different signal to noise profile: the most probable peaks had the highest and most narrow peak of signal, while less probable peaks had progressively lower and broader signal domains (Figure 5C). The signal profile of a peak is thus an indicator of its probability of detection.

We integrated RNA-seq of HCT116^28,29^ to determine correlation between peak detection probability and gene transcription. Hypergeometric tests revealed that overall, ICEBERG peaks were associated to transcribed genes (59% of peak associated genes expressed: 1.27-fold enrichment, p-value 9.54 e^-242^, transcripts per million (TPM) > 5, peaks annotated to genes using GREAT) (Figure 5D, left). This association increased when considering only peaks occurring in promoters, where peak-gene annotation is more precise (70.5% of peak associated genes expressed, 1.51-fold enriched, p-value 3.95 e^-186^) (Figure 5D, center). The rare peaks were equally associated to occurring transcription as shown by both percentage of expressed peak annotated genes, and their level of expression (Figure 5D, right), indicating that they are part of a biological pathway active in HCT116.

Genomic annotations of peak positions revealed that the high probability peaks were typically associated with intergenic regions and poorly associated with promoters (51% intergenic, 4% promoters) confirming what has been previously shown for TCF/LEF and β-catenin^26^. However, the rarest peaks were statistically more enriched in promoters, and less so for intergenic regions (29% intergenic, 28% promoters: 4.6-fold enrichment, logP^-4484^) (Figure 5E). Accordingly, the rarest peaks were more often marked by H3K4me3 than the high probability peaks (51.6% vs. 39.2%) (Figure 5F). The rarest peaks also appeared to be more often associated with non-Tn5-accessible chromatin within a cell population (ENCODE ATAC-seq for HCT116, IDR threshold peaks)^30^ (Figure 5G). This could reflect that these rare β-catenin binding instances are associated with a smaller fraction of cells in which the chromatin might be accessible, but not detectable by bulk level ATAC-sequencing.

### Peak probability identified by ICEBERG predicts cis-regulatory element interactions

We have shown that β-catenin peaks can be divided into probability groups that are characterized by different features such as signal intensity, promoter association and chromatin accessibility. This raises the prediction that β-catenin binding event probability could be associated with differential functionally relevant regulatory events. To test our prediction, we focused on active cis-regulatory three-dimensional chromatin interactions by employing published HiChIP of H3K27ac in HCT116^31^. This assay uncovers distant regions that are engaging in gene regulation events and that are marked by the open chromatin mark often found in active enhancer regions^31^. We focused on H3K27ac mediated loops that overlap with a β-catenin ICEBERG peak in at least one anchor and assessed the correlation between β-catenin peak detection frequency and 1) number of loops associated to each probability group, 2) loop strength in terms of counts and 3) loop size in terms of linear genomic distance between the two anchors. We observed a strong correlation between peak probability and number of associated loops: high probability peaks were connected on average to ∼6 loops, while the lowest probability peaks were connected to ∼1 (Figure 6A, left). Moreover, both loop strength and size showed a similar pattern, with higher probability peaks associated with stronger and larger loops (Figure 6A, center and right).

**Figure 6.**
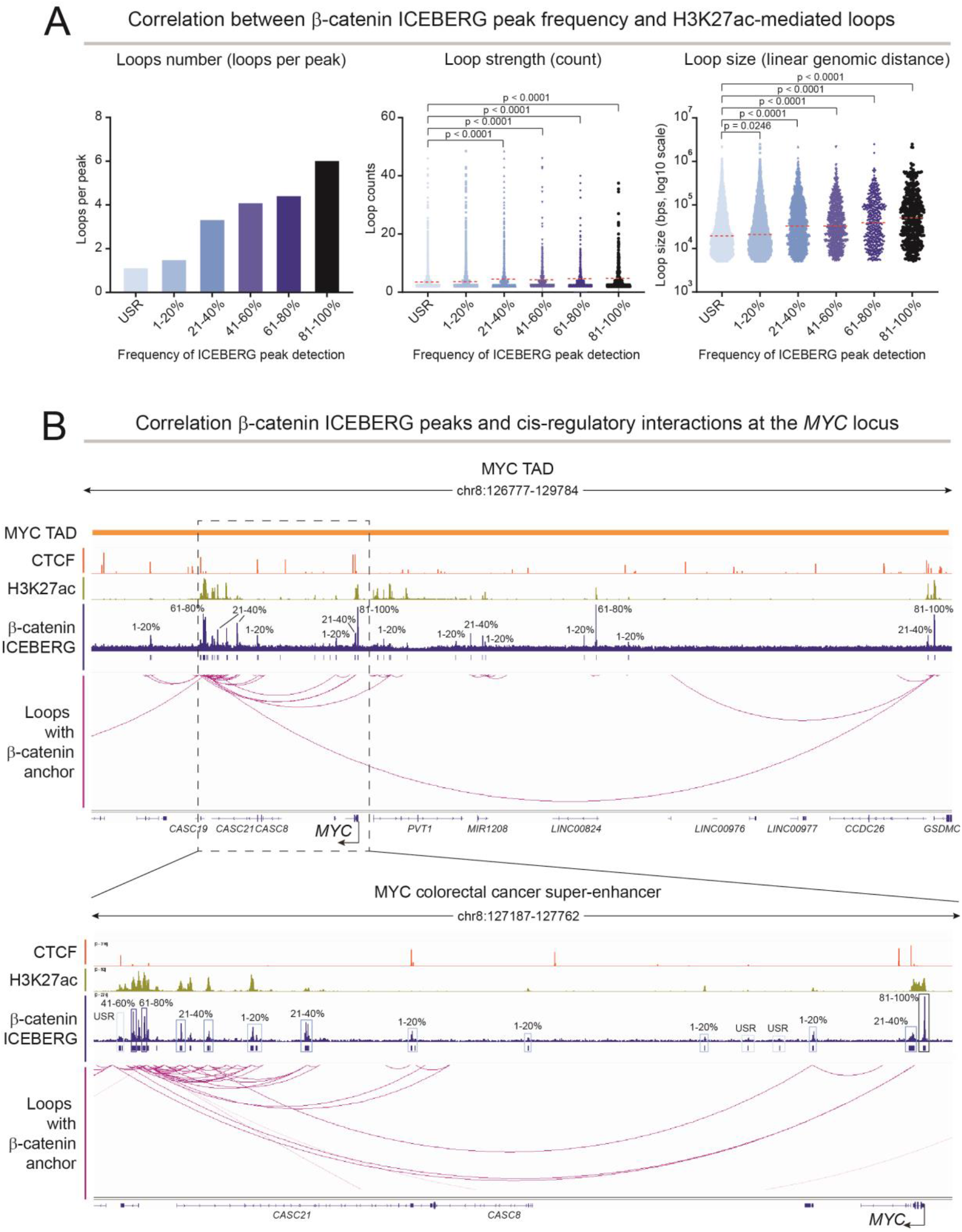
ICEBERG peak probability is correlated with cis-regulatory element activity. **A**. Left: bar graph representing average number of H3K27ac mediated chromatin loops as seen by HiChIP, per β-catenin peak in the different probability groups. Increasing probability of detection correlates with increasing number of chromatin loops with at least one anchor in the β-catenin peak. Middle: Graphs of loop strength distribution (measured by average count) for loops associated with different β-catenin peak probability groups. The strength of loop interactions with the most probable peaks is higher than for the least probable peaks. Right: graphs of loop size distribution (measured by linear distance in kb, log transformed) for loops associated with β-catenin peaks. Higher probability peaks are associated with larger loops. **B**. Top: representation of the *MYC* topologically associating domain (TAD), showing CTCF ChIP-seq, H3K27ac ChIP-seq, the β-catenin ICEBERG dataset, and β-catenin anchored H3K27ac HiChIP loops. Bottom: zoom-in on the *MYC* gene body and the colorectal cancer super enhancer region.

We turned our attention to the highly dynamic, cancer relevant and well-studied *MYC* locus, and its super enhancer^32^. ICEBERG unearthed a plethora of β-catenin binding events within the *MYC* topologically associated domain, ranging in probability of detection (Figure 6B). The vast majority of peaks are associated to a H3K27ac loop, hence are functionally active. Within this locus, even low probability and undetectable in single replicate peaks (USR) are connected to loops. Our analysis highlights that while most β-catenin and H3K27ac loops are within the colorectal cancer *MYC* super-enhancer region and *MYC* gene body (Figure 6C), others can be found further upstream, connecting the super-enhancer over 2 Mb distance with a high probability β-catenin peak near the TAD boundary, implying that different β-catenin bound loci may have different functions.

### ICEBERG identifies cancer relevant, rare β-catenin direct targets

We set out to discover which are the rarest β-catenin ICEBERG binding events. To do this, we started from the 6708 USR ICEBERG peaks (undetectable in single replicates, q < 0.05 as per Figure 5B), and then progressively relaxed our stringency thresholds. 1897 ICEBERG peaks remained undetectable in single replicates at lower stringency (p < 0.001), and of these, 1713 ICEBERG peaks were called less than 5 times at the lowest stringency (p < 0.01). Finally, we defined the 524 ICEBERG peaks called in only one individual replicate with the lowest stringency (p < 0.01) as the rarest β-catenin binding events (Figure 7A). This group of rare events constitutes a new discovery made possible by ICEBERG. In fact, they can be identified in single replicates only by loosening the stringency to a point where false positives become overwhelming in number (pre-cision < 0.2, Figure 4B). Consistently, even a published ChIP-seq replicate for β-catenin in HCT116, which makes use of crosslinking, large number of cells, and high sequencing depth, only identified 7/524^33^.

**Figure 7.**
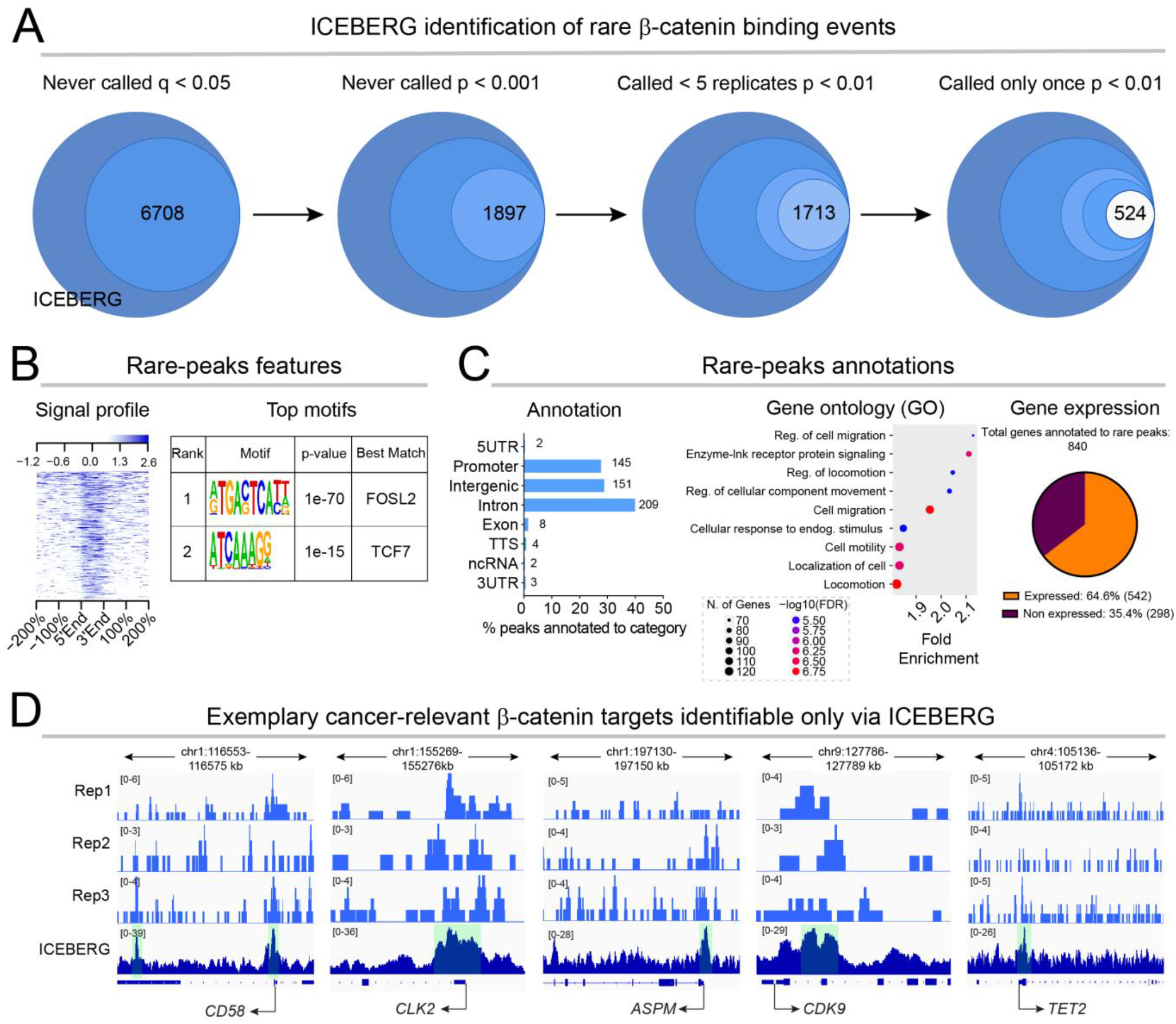
ICEBERG identifies rare β-catenin targets relevant for colorectal cancer. **A**. Schematic representation of the definition of the 542 rarest β-catenin peaks. **B**. Left: signal intensity plot of the ICEBERG β-catenin dataset over IgG negative control within the rarest peak regions, showing up to 2.6-fold enrichment over control. Right: HOMER *de novo* motifs show AP1 and TCF/LEF as the top enrichment motifs within the rarest peaks, confirming their credibility. **C**. Left: bar graph showing genomic region annotation, displaying that the rarest peaks fall within mostly promoter, intergenic and intronic regions. Center: Dot plot showing the top 10 enriched gene ontology biological processes for the rarest peak-annotated genes (cutoff FDR < 0.05). The x-axis shows fold enrichment, dot size is dependent on gene number, and color represents - log10(FDR). Many terms involve motility, movement, and migration. Right: pie graph showing the expression status of the rarest peak annotated genes, which are similarly expressed to the ICEBERG dataset as a whole (See Figure 5D). **D**. Visualization of the β-catenin ICEBERG dataset and example single replicates on the notable *CD58, CLK2, ASPM, CDK9* and *TET2* loci.

Despite being rare events, they displayed up to 2.6-fold signal enrichment over the control and were characterized by the presence of AP1 and TCF/LEF family motifs as the most enriched (Figure 7B). They displayed one of the highest enrichments for promoters that we have seen across groups of β-catenin targets (28% in promoters) (Figure 7C, left). We annotated the 524 peaks to 840 genes using GREAT and performed gene ontology analysis to determine whether these rare β-catenin binding events could be associated to specific biological processes. Terms related to cell motility and migration were the most enriched (FDR < 0.05), indicating that they could be highly relevant for cancer invasiveness and metastasis (Figure 7C, center). 64.6% of the genes annotated to peaks were expressed in HCT116 (> 5 TPM) (Figure 7C, right). Among the highest expressed are notable, cancer relevant players which have never been previously identified as direct targets of β-catenin (Figure 7D). *CD58*, for example, is a surface marker of CRC tumor-initiating stem cells that has previously been found to be expressed by only ∼ 4% of cells in CRC cell lines, correlating well with the observed rarity of the β-catenin peak, yet is capable of enhancing self-renewal and promoting epithelial to mesenchymal transition^34^. *CLK2*, CDC-like kinase 2, is known to be upregulated in CRC and is associated with poor survival. It is involved in splicing of Wnt pathway genes, and Wnt has been shown to be responsible for the CLK2 induced cancer progression^35,36^. CLK2 inhibitors are now being tested in clinical trials to treat Wnt-driven CRC^37^. *ASPM* has been identified as a biomarker for CRC and is upregulated in CRC tissues, functioning to promote proliferation and inhibit apoptosis, affecting the cell cycle^38^. ASPM levels are positively correlated with β-catenin levels^39^. *CDK9*, Cyclin-dependent kinase 9 plays a role in transcriptional elongation and has been shown to be a potential drug target in CRC^40^. *TET2*, ten-eleven translocation 2, acts as a tumor-suppressor in many types of cancer via its enzymatic DNA demethylation activity^41^. This warrants screening of all the ICEBERG hits for their developmental or disease relevance.

## Discussion

Current technologies such as the popular ChIP-seq and CUT&RUN provide experimental means to detect protein-DNA interactions genome-wide. However, as they rely on NGS, they are intrinsically noisy and pose the challenge of drawing arbitrary thresholds, which when relaxed can increase discovery with the cost of including false signal, and when stringent, become more precise at the expense of excluding real signal^15,18^. Here, we attempt to address this problem using CUT&RUN, though this issue is central to the signal detection theory and is not limited to these types of technologies.

As scientists we typically prefer not to be wrong; thus, a common philosophical stance in molecular biology is to prefer false negatives over false positives, which increases our confidence in the reliability of the findings but limits the breadth of the experimental outcome. Here, we propose that this might not always be the best strategy. In the clinics, for example, it is arguably preferable to sustain the cost of investigating minor suspicions, rather than leaving a disease undiagnosed. This is not only an intellectual exercise, as the discoveries concerning binding of transcription factors are now employed for CRISPR-based mutagenesis strategies in clinical trials^42^.

ICEBERG attempts to prevent the need for compromise between false positives and false negatives. By increasing the capture of signal enrichment through replicate aggregation until saturation of peak discovery, ICEBERG is able to average out background and expose reproducible yet typically undetectable signal enrichment. When applied to β-catenin, a notoriously difficult target, ICEBERG built with 25 replicates uncovers more than 93% of all targets according to the fitted function. Moreover, our calculations indicate that each additional replicate after 20 contributes to a proportionally small group of additional peaks. ICEBERG can therefore be adapted to understand how many replicates are needed for a given factor. We believe that this will depend on biophysical properties and mechanism of action of each factor.

We leveraged our datasets to benchmark common experimental strategies and peak calling pipelines, and provide estimates of how these perform for different targets. Our results indicate that duplicates of a stable chromatin mark like H3K4me3 are sufficient to identify a significant portion of the ICEBERG with high precision. However, β-catenin, presumably as other dynamic gene regulators, requires more datasets to approach the totality of binding events. While ICEBERG might seem demanding, note that we used a total of 6,250,000 cells (250,000 per replicate), 3.5 μg of antibody (0.14 μg per replicate), and 125 million reads (5 million per replicate) to produce it. These numbers are grossly comparable to the requirements of a single ChIP-seq setup. Working with HCT116, we did not need to scale down in terms of cell number, but CUT&RUN is routinely successfully performed on < 10,000 cells^43^, rendering ICEBERG applicable virtually to every tissue or cell type, and friendly to scalability and automatization.

One could consider that a limitation of the ICEBERG is the difficulty in identifying an orthogonal technology to validate it. Historically, CUT&RUN has been benchmarked against a single or two replicates of ChIP-seq. However, ChIP-seq itself suffers from the same downfalls as CUT&RUN and has a similarly low overlap between replicates^15,44^. Limited replicates of ChIP-seq would thus constitute an underpowered comparison to a CUT&RUN ICEBERG; on the other hand, we posit that ChIP-seq experiments would require their own ICEBERG. Similarly, this reasoning could be applicable to other high throughput NGS-based approaches.

ICEBERG does more than just identify all binding events – it also associates them to a spectrum of detection probabilities. Instead of a binary (yes or no) view of binding events, each chromosomal position is associated with a probability of being bound by a given factor. Regions with the highest signal in the ICEBERG are associated with a high probability of detection as they are identified in all or nearly all individual replicates. On the other side of the spectrum, regions with significant yet lower signal in the ICEBERG datasets are rarely detected in single replicates. We suggest two non-mutually exclusive explanations for this: 1) detection probability reflects the fraction of cells in which the event occurs, and 2) the detection probability is proportional to the actual time a factor remains associated to that locus. Single-cell approaches may provide the evidence for the first explanation^8^, while biophysical readouts could be used to assess the second^45^.

Finally, applying ICEBERG to β-catenin in CRC cells provided important biological insights. First, it produced an exhaustive map of its binding pattern, which is all the more critical as β-catenin has been a historically challenging protein to target with these types of assays. Second, ICEBERG revealed more than 500 cancer relevant binding instances never previously identified, underlying so far neglected regulatory events of the Wnt/β-catenin pathway. Among these new target genes, are some that are now undergoing clinical trials as drug targets to treat CRC and for which evidence was presented indicating that they are genetically downstream of Wnt signaling^35,37^. We consider this as an important validation of our approach. As even individual enhancers identified as binding sites of transcriptional modulators by CUT&RUN are now being targeted by CRISPR/Cas9-mediated deletion in clinical trials^42,46^, our identification of rare and cancer-relevant regulatory sites demands urgent investigation.

## Acknowledgments

This work was supported by Grants to C.C. from Cancerfonden (CAN 2018/542 and 21 1572 Pj), the Swedish Research Council, Vetenskapsrådet (2021–03075), and Linköping University. C.C. is a Wallenberg Molecular Medicine (WCMM) fellow and receives generous financial support from the Knut and Alice Wallenberg Foundation. Computations and data handling were enabled by resources provided by the Swedish National Infrastructure for Computing (SNIC) at [SNIC CENTRE] partially funded by the Swedish Research Council through grant agreement no. 2018-05973.

## Author contributions

A.N., P.P., G.Z., and C.C. conceived the project. A.N., P.P., and G.Z. performed the experiments. A.N. and P.P. performed the formal analysis and prepared figures. C.C. supervised the research team and provided financial support for the study. A.N., P.P., and C.C. wrote the manuscript “with six hands”. All authors reviewed and commented on the final manuscript.

## Availability of Data and Materials

Publicly available datasets used in this study can be downloaded according to their accession information. CUT&RUN raw data and ICEBERG tracks for visualization have been deposited at Array Express, accession number E-MTAB-13143. ICEBERG peak sets, annotated for probability group, and annotated genes are provided as supplementary data. Additional processed files and scripts are available upon request.

## Methods

### Cell culture

HCT116 human colorectal cancer cells were cultured in a humidified 37 °C incubator in 5% CO_2_. Culture medium consisted of high glucose Dulbecco’s Modified Eagle Medium (Cat. #41965039, Gibco) with 10 % bovine calf serum (Cat. #1233C, Sigma-Aldrich) and 1X Penicillin-Streptomycin (Cat. #15140148, Gibco).

### CUT&RUN LoV-U

CUT&RUN LoV-U was performed as described in Zambanini et al.^7^. 250,000 cells/sample were harvested from culture flasks by incubation with Trypsin-EDTA (Cat. # 25200056, Gibco) for 5 minutes. The trypsin was quenched with culture media and the cells were washed twice with DPBS (Cat. #14190094, Thermo Fisher Scientific). Nuclear extraction was performed by three washes Nuclear Extraction (NE) buffer (HEPES-KOH pH-8.2 [20 mM], KCl [10 mM], Spermidine [0.5 mM], IGEPAL [0.05%], Glycerol [20%], Roche Complete Protease Inhibitor EDTA-Free). The nuclei were resuspended after the final wash in 40 μl NE per sample and bound to 10 μl Magnetic ConA Agarose beads equilibrated in binding buffer (HEPES pH 7.5 [20 mM], KCl [10 mM], CaCl_2_ [1 mM], MnCl_2_ [1 mM]) as described in Meers and colleagues^43^. Bead binding proceeded on a rotator for 15 min, after which nuclei and beads were resuspended in 200 μl wash buffer per sample and split into PCR tubes. Beads were collected on the magnet and then resuspended in 200 μl EDTA wash buffer (wash buffer with EDTA [0.2 mM]) and incubated RT for 5 min. Samples were then directly resuspended in antibody buffer (wash buffer with antibody 1:100) and incubated ON at 4 °C on a rotator. Antibodies used included anti-β-catenin ABIN2855042, anti-H3K4me3 ABIN6971977 and anti-rabbit IgG ABIN101961. After ON incubation samples were washed 5 times and re-suspended in 200 μl of pAG-MNase buffer (wash buffer with pAG-MNase 0.6 μg/ml) and incubated for 45 min at 4 °C on a rotator. pAG/MNase was a gift from Steven Henikoff (Addgene plasmid #123461; http://n2t.net/addgene:123461;RRID: Addgene_123461) and was expressed and purified in house as previously described^43^. Next, samples were washed five times and during the last wash equilibrated 5 min in wet ice. Samples were resuspended in 200 μl ice cold wash buffer containing 2 mM CaCl_2_, and digestion was allowed for proceed for exactly 30 min in wet ice. The digestion buffer was removed and transferred to tubes containing 1.5 μl of 0.5 M EDTA and 1.5 μl of 0.5 M EGTA and mixed well. The reaction was stopped on the beads with 47 μl of 1X Urea STOP buffer (NaCl [100 mM], EDTA [2 mM], EGTA [2 mM], IGEPAL [0.5%], Urea [8.8 M]) and the samples were incubated 1 hr at 4°C for elution. Beads were collected on the magnet and liquid transferred to the PCR tubes containing the digestion buffer. DNA purification was done with two rounds of bead purifications according to manufacturer’s directions, using Mag-Bind TotalPure NGS beads (Cat. #M1327, Omega Bio-Tek) at 2X (200 μl first round and 40 μl second round), and then finally re-suspended in 20 μl Tris-HCl pH 7.5.

### Library preparation and sequencing

Library preparation was done using the KAPA Hyper Prep Kit for Illumina sequencing (Cat. #KK8504, KAPA Biosystems) following the manufacturer’s guidelines with modifications. 0.4X volume reactions were used for End repair and A-tailing steps, starting with 20 μl CUT&RUN DNA. The thermocycler conditions were 12 °C for 15 min, 37 °C for 15 min and 58 °C for 25 min. Adapter ligation was also performed in 0.4X volume reactions. KAPA Dual Indexed adapters were used at 0.15 μM. A DNA cleanup was performed after ligation with Mag-Bind TotalPure NGS beads at 1.2X the sample volume, and DNA was resuspended in 10 mM Tris-HCl pH 7.5. Library amplification steps were performed in 0.5X volume reactions. The cycling conditions were: 45 sec initial denaturation at 98 °C, 15 sec denaturation at 98 °C, 10 sec annealing/elongation at 60 °C, 1 min final extension at 72 °C, hold at 4 °C, for a total of 13 cycles. A post-amplification DNA purification was performed with beads at 1.2X sample volume. Libraries were then size-selected using the E-Gel EX 2% agarose gel (Cat. # G402022, Invitrogen) and the E-Gel Power Snap Electrophoresis System (Invitrogen). Run time was set to 10 minutes and DNA between 150 and 500 bp was excised from the gel and purified using the QIAquick Gel Extraction Kit (Cat. #28706X4, Qiagen) according to the manufacturer’s instructions. Libraries were quantified using Qubit (Thermo Scientific) and the high sensitivity DNA kit (Cat #Q32854, Thermo Scientific). Libraries were pooled according to nM and sequenced with 36 bp pair-end reads on the NextSeq 550 (Illumina) using the Illumina NextSeq 500/550 High Output Kit v2.5 (75 cycles) (Cat. #20024906, Illumina). Target depth was 5 – 10 million reads per sample.

### Replicate Quality Control and Alignment

Fastqc (version 0.11.9)^47^ was used to assess initial read quality and duplication rates. Reads were trimmed using bbmap bbduk (version 38.18)^48^ to remove adapters, known artifacts, poly AT and TA repeats, and poly G and C repeats. The trimmed reads were aligned to the hg38 genome with bowtie2 (version 2.4.5)^49^ with the options –local –very-sensitive-local –no-unal –no-mixed -no-discordant –phred33 – dovetail -I 0 -X 500. Samtools (version 1.11) ^50^ was used to convert sam to bam files, fix improperly paired mates, and remove duplicates. BEDTools (version 2.30.0)^51^ was used to remove reads mapped to the CUT&RUN hg38 blacklist^13^. Final read counts of usable fragments were determined from filtered and deduplicated bam files. Bedgraphs were created with BEDTools (version 2.23.0)^51^ genomecov on pair-end mode. Quality control correlation plots and PCAs were performed with deepTools (version 3.5.1-0)^52^. Bedgraphs were visualized in IGV^53^.

### Peak Calling and Decay Curves

The 10 IgG negative controls were merged to create the aggregate IgG (approximately 70 million reads). The aggregate IgG was downsampled to 5 million reads for peak calling of individual replicate datasets. Initial peak calling was performed with MACS2 (version 2.2.6)^10^ with the options - f BAMPE --keep-dup all and -q 5e-2 with the downsampled IgG as the control. Intervene (version 0.6.4)^54^ was used to create Venn diagrams. BedSect^55^ was used to overlap all replicates, and the resulting analysis matrix files were used to create decay curves. The peaks were determined using awk to extract the exact regions called in at least *x* datasets, and then merging neighboring regions with BEDTools and calculating the number of unique regions. Peaks called in exactly *x* datasets were calculated by subtracting the *x+1* peaks from the *x* peaks. Curves were plotted in R.

## ICEBERG

The maximum number of usable fragments was determined by the smallest replicate to be 4 million for β-catenin and 5 million for H3K4me3, and each replicate was downsampled to this depth 3 different times; reads were shuffled before each downsampling. The downsampled replicates were merged one by one for each of the three splits to create three different aggregate datasets, containing 100 million reads for β-catenin and 50 million reads for H3K4me3, coming from 25 replicates and 10 replicates, respectively. Each of the three aggregates were separately peak called with MACS2 as described above, against the aggregate IgG dataset. Between each replicate addition, peaks were also called in the same way and the number of peaks were used to generate the ICEBERG build curves. This was done 5 times with the replicates in different random orders. The relationship between the number of aggregated replicates and called peaks was modeled by fitting a series of polynomial regressions of increasing order (up to 5^th^) in R (version 4.2.2, lm function). The function with the highest R^2^ value was selected. b-catenin ICEBERG aggregation was modeled by a third order / cubic regression, [0.08184^*^x^3-20.94564^*^x^2+1036.53907^*^x-812.39289], while H3K4me3 ICEBERG aggregation was modeled by the following fifth order regression: [-0.4687^*^x^5+10.3849^*^x^4-74.7615^*^x^3+176.2564^*^x^2+370.7989^*^x+14700.3867]. The first derivative was then calculated for both functions and plotted. Peaks called in at least 2 of the 3 aggregates were determined with BEDTools, and then further filtered to remove peaks not called in at least one individual replicate when called with MACS2 -p 1e-2, to finally generate the final set of ICEBERG peaks. The ICEBERG peak sets were overlapped with the peak data for the decay curve of the replicates in order to create the ICEBERG peak decay curve and define the probability groups as explained in the results.

### Benchmarking of Replicates and Peak Callers

I-FrIPs were calculated as (reads within ICEBERG peaks)/(total mapped reads). For the benchmarking, MACS2 was used as described above, and also with -p 1e-2 and -p 1e-3 as thresholds. SEACR (version 1.3)^11^ was used with the settings norm relaxed against IgG, norm stringent against IgG, and 0.001 stringent. For the 0.001 peaks, the same settings were applied to the IgG downsampled negative control, and peaks called in the negative control were subtracted from each replicate peak set with BEDTools subtract downstream. Precision was determined by (peaks within ICEBERG set)/(total called peaks), and recall as (peaks within ICEBERG/total ICEBERG peaks). F1 scores were calculated by the equation F1 = 2^*^(Precision^*^Recall)/(Precision+Recall). Duplicate peak sets were obtained by using BEDtools intersect to overlap the peak sets of all the possible replicate combinations. For triplicates, random datasets were overlapped, and both the 2 of 3 and 3 of 3 peak sets were generated with BEDtools. Redundant results (those containing the same dataset multiple times) were removed.

### Downstream Analyses and Graphs

Signal intensity plots and average profiles were created using ngs.plot (version 2.63)^56^ with options -G hg38 -R bed -N 2 -SC global -IN 0 for peak regions, and with the options -G hg38 -R tss -L 1500 – SC global for the whole genome TSS H3K4me3 plot. Genomic region annotation was done with HOMER (version 4.11)^57^ annotatePeaks on default settings, using –annStats to extract annotation statistics. Genes assigned to promoters were extracted from HOMER results. Motif analysis was done with HOMER with the -size given parameter to find *de novo* motifs. Peak annotation to genes (not only those in promoters) was done using GREAT (version 4.0.4)^58^ on default settings. Gene ontology was performed for GO biological processes using ShinyGO^59^, results were cutoff at FDR of 0.05.

### Analysis of Published CUT&RUN, RNA-seq, ATAC-seq, HiChIP, and ChIP Data

CUT&RUN datasets were downloaded for the following factors: SALL4^60^; Ikaros and H3K4me3^61^; GATA1^62^; H3K27ac^63^; H3K9me3^64^; SOX2^65^; HDAC2^66^; β-catenin, H3K4me2, HDAC1, CBP^7^. Peak sets, exactly as they appeared in the publications, were overlapped with BEDTools intersect. Concordant peaks were determined as peaks in the overlap divided by the total peaks called, expressed in percentage.

Baseline RNA-seq data in the form of transcripts per million (TPM) for HCT116 were downloaded from the studies in reference here^28,29^. Transcript IDs were converted to gene names, and the lists were filtered for expressed genes as defined by a TPM of > 5. Only genes expressed in both datasets were kept. The resulting list of expressed genes was compared with peak annotated gene sets generated as described above. Hypergeometric tests were performed in R with phyper.

ATAC-seq data for HCT116 from the ENCODE project was used^30^. IDR thresholded peaks were overlapped using BEDTools with the ICEBERG peaks from each group to define the peak regions as open or closed chromatin.

ChIP-seq peaks of β-catenin in HCT116 were downloaded from Bottomly and colleagues^33^ and lifted over to the hg38 genome using UCSC liftover^67^. Fold-change over control bigwig files mapped to hg38 were downloaded for H3K27ac conducted in HCT116^68^, and for CTCF conducted in HCT116^69^, performed by the ENCODE consortium.

HiChIP of H3K27ac in HCT116 data was performed by Chen and colleagues^31^. bedpe files containing unfiltered loops downstream of HiChipper were downloaded from GEO GSE173699. Loops were filtered, keeping those present in both biological replicates, with a size greater than 5 kb, and a count average of 2 or greater. UCSC liftover was used to convert hg19 to hg38 coordinates, and then loops were overlapped with β-catenin ICEBERG peaks using BEDTools pairtobed, retaining only loops with one or more anchors overlapping a β-catenin peak. β-catenin anchored loops were then separated into groups based on peak probability, and the number of loops was divided by number of peaks to obtain average loops/peak. Loop strength was determined for each loop by the average loop count of the two replicates, and loop size was determined by the linear genomic distance between the start of the first anchor and the end of the second anchor. Log10 of the loop size was used for the plots. Statistical testing was done using non-parametric Kruskal-Wallis test with pairwise comparisons and multiple testing corrections, using GraphPad Prism.

